# Acute lymphoblastic leukemia cells image analysis with deep bagging ensemble learning

**DOI:** 10.1101/580852

**Authors:** Ying Liu, Feixiao Long

**Author notes:** Corresponding author: Feixiao Long.

## Abstract

Acute lymphoblastic leukemia (ALL) is a blood cancer which leads 111,000 depth globally in 2015. Recently, diagnosing ALL often involves the microscopic image analysis with the help of deep learning (DL) techniques. However, as most medical related problems, deficiency training samples and minor visual difference between ALL and normal cells make the image analysis task quite challenging. Herein, an augmented image enhanced bagging ensemble learning with elaborately designed training subsets were proposed to tackle above challenges. The weighted *F*_1_-scores of preliminary test set and final test are 0.84 and 0.88 respectively employing our ensemble model predictions and ranked within top 10% in ISBI-2019 Classification of Normal vs. Malignant White Blood Cancer Cells contest. Our results preliminarily show the efficacy and accuracy of employing DL based techniques in ALL cells image analysis.

## 1 Introduction

Acute lymphoblastic leukemia (ALL) is a blood cell cancer caused by the development of immature lymphocytes. About 876,000 people suffer ALL globally in 2015 and lead to 111,000 death according to the data shown in [1]. The treatment of ALL normally includes the chemotherapy which may last for several years. Typically, diagnosis of ALL is through blood tests and bone marrow biopsy etc. Among several tests, microscopic image based inspection is one of the promising and easy implemented methods, especially compared with expensive bio-chemical based inspections such as cell flow cytometry. The method through microscopic images to detect and classify various cells (not limited to ALL cells) can be viewed as a kind of well-defined and investigated computer vision (CV) problems. Recently, with developing of deep learning (DL) based techniques, researches have investigated the potential use of applying DL methods in cell image classification problems. Compared with conventional hand-craft features CV based solutions, DL based techniques have the advantages with automatically selecting representative image features of cells [12] and reduce the difficulty of solving ALL cells classification through a series of standard DL procedures.

For example, Shafique etc. [9] employed upgraded Alex Net to differentiate ALL cells as well as their subtypes. Rehman A. etc. [7] also employed convolutional neural network (CNN) to classify ALL cells between normal cells. Besides, the authors compared the DL based methods to conventional machine learning (ML) approaches, such as k-nearest neighbours (k-NN), support vector machine (SVM) etc. Wang etc. [13] proposed a marker-based learning vector quantization neural network to detect and classify ALL cells.

However, although multiple researchers have shown the efficiency of applying DL based techniques in ALL cells image analysis, detecting ALL cells based on microscopic images would still be very challenging due to multiple reasons. First, in most medical related object detection and classification problems, it requires professional medical knowledge or experiences to correctly label the objects. This often results in the deficiency training samples compared to nature image processing tasks. Second, to medical related datasets, it is common to observe that the imbalance between the number of positive and negative or different categorical samples due to the nature of disease or physiological situations. For example, the number of ALL cells is approximately twice than the number of healthy cells (denoted as HEM) in the dataset (https://competitions.codalab.org/competitions/20429#learn_the_details-data-description) employed in the experiment. From the aspect of machine learning, intuitively the imbalance issue will push the model to ‘learn’ representative features from majority samples only and neglect minority ones without proper settings during training. Therefore, careful oversampling or undersampling of training sets as well as other strategies have to be performed in order to reduce the influences of imbalance datasets. Last, as Fig. 1 shown, it is not easy to differentiate ALL and HEM cells image visually without professional experiences. In other words, the ALL cells discrimination problem suffers relative small inter-class similarity, which increases the difficulty of automatic detection.

**Fig. 1.**
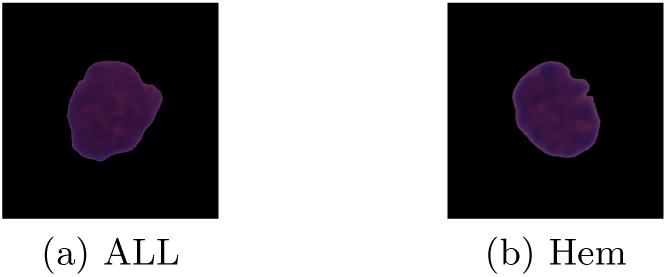
Close visual similarity between ALL and HEM cells

In order to address above issues, transfer learning [11] was employed to deal with relative deficiency training samples. Specifically, pre-trained Inception ResNet V2 [10] using ImageNet (http://www.image-net.org) was employed as backbone net. Besides, a two-stage augmented image enhanced bagging ensemble training strategy [6] was employed to reduce the influences of imbalance dataset. Two subsets (see Sec. 2.3) were carefully designed for training backbone nets followed the idea shown in [6] in the first step training. Inspired by [15], two separately trained models were ‘combined’ together and re-trained with all training samples in order to further boost the overall prediction performance. Note that for simplicity, binary cross entropy loss rather than synergic loss was employed during the training of ‘combined’ model.

The manuscript is arranged as following. In Methods section, our carefully designed training subsets will be described in details. Moreover, the training settings and strategy will be discussed and introduced. In Results section, the accuracy of the model will be illustrated. Finally, several issues of employing DL techniques in medical related problems in the future will be discussed.

## 2 Methods

### 2.1 Details of initial dataset

As described in the contest (https://competitions.codalab.org/competitions/20429#learn_the_details) [2, 4], the initial dataset consists of 76 subjects, among which 47 subjects suffer ALL and the rest are healthy. Totally, there are 10,661 cells, in which 7,272 cells are classified as ALL and 3,389 cells are classified as HEM. Although there is only one cell in each training image, the image itself is not the convex hull of the cell and thus more black background than necessary is contained in the image. Herein segmentation of cell to get the convex hull through common image processing techniques was applied to the image (see Sec. 2.2) in order to get rid of most black background. Staining and illumination error were presented in the images due to image capture with real clinical situations although reduced by stain normalization techniques as shown in [3,5]. The initial dataset was then split into original training set (see Sec. 2.3 for more details on generating final training set A and B) and validation set with proportion 7:3 as conventional settings in DL.

The preliminary test set consists of 30 subjects, in which 15 subjects are labeled as ALL and the rest are HEM. Similar cell segmentation and stain normalization techniques as described above were employed. In total, the preliminary test set consists of 1,867 cells, among which 1,219 cells were classified as ALL and 648 cells were HEM.

### 2.2 Images pre-processing

As shown in [14], the authors employed a segmentation network first to segment the cell area from the image, making the classification network focus on representative and specific cell region [15] without being distracted by the background. Inspired by this idea, the cell images were first pre-processed through a pipeline of image processing techniques to filter out most black background. Briefly, the RGB image was transferred to gray scale one and the region where the cells locate was binarized using the threshold estimated by Otsu’s method followed by the erosion operation. The convex hull of the binarized region was then calculated and the RGB patch containing the cell in the middle was extracted accordingly. Finally the image was resized to 299 × 299 × 3 based on the input requirement of the employed backbone Inception ResNet [10] model.

### 2.3 Training dataset generation

Considering the number of ALL cells is approximately double of the number of HEM cells, we decided to adopt modified bagging ensemble training strategy [6] (briefly speaking two models employed, with each model learns different partial image features from majority classes but same features from minority classes intuitively) to deal with this imbalance dataset and train the backbone models separately at the first stage training (see Sec. 2.5 for more details). The ALL cell samples were divided into two equally number subsets (named as ‘All-A’ and ‘All-B’ respectively, see Fig. 2). Moreover, randomly chosen one or combination of several common image augmentation techniques, such as rotation, horizontal or vertical flip, Gaussian noise addition as well as contrast and color channel adjustment etc., was applied to ALL cell subsets and HEM cell set for increasing the number of training samples and reducing over-fitting of the model. As shown in Fig. 2, the training set A consists of ALL samples from subset All-A and the augmentation images form subset All-B. In contrast, training set B consists of ALL samples from subset All-B and augmentation images from subset All-A. The HEM cells as well as their augmentation part were employed in both training sets (A and B). Intuitively, each backbone model will ‘learn’ the same HEM cell features but ‘mixed’ ALL cell features from different compositions of All-A, All-B and their image augmentation counterparts.

**Fig. 2.**
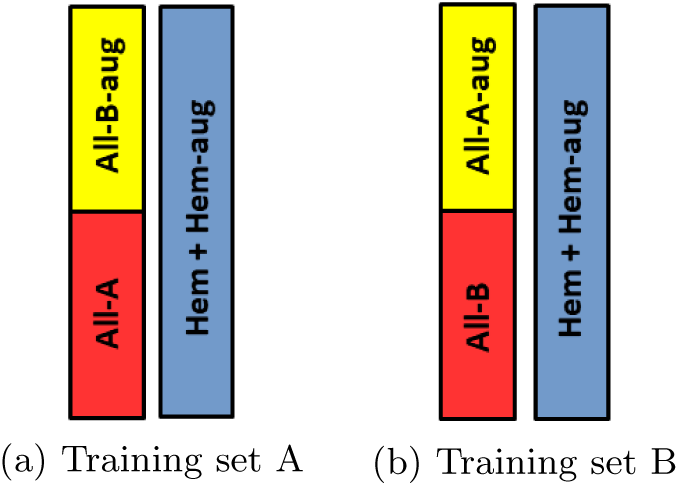
Composition of training sets

### 2.4 Training strategy and ‘combined’ model architecture

Two stage training strategy was employed in our experiments as described below for further increasing the overall model performance. For the first stage training, two Inception ResNets (denoted as model-A and model-B) with pre-trained ImageNet weights as initial were employed to be trained on the training set A and B respectively. For the second stage training, models trained in the first stage were ‘combined’ (See Fig. 3) and fine tuned using all the ALL and HEM cells in the training set. In details, the pre-trained Inception ResNets in the first stage were employed as image features extractor. The outputs from penultimate layers (global average pooling layer, 1536-dim vector) from each Inception ResNet were concatenated after the batch normalization operation (not shown in Fig. 2 for simplicity) for reducing the influences of possible large magnitude differences between two output feature vectors from model-A and model-B respectively. Next, two fully connected (FC, 1024 neurons each) layers with dropout operation (dropout rate = 0.6) were added to increase the prediction accuracy of the ‘combined’ model. The output layer of the ‘combined’ model was another FC layer with softmax activation function conventionally. This model, as its name shown, combines learned partial representative features from two ALL subsets and will further increase the prediction accuracy.

**Fig. 3.**
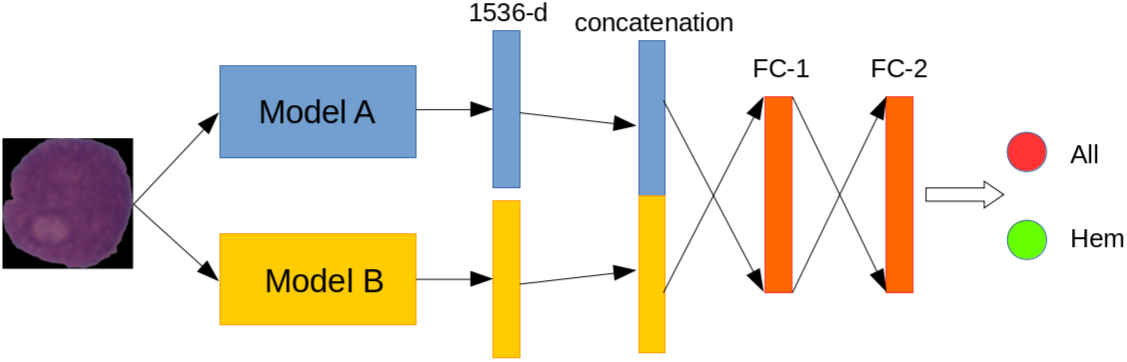
Structure of the ‘combined’ model

### 2.5 Training settings

During training, RMSprop optimization [8] with all the settings default was employed. The total epochs were set to 50 empirically. In order to reduce the over-fitting and increase the performances of all models (model-A, model-B and ‘combined’ one), several common techniques were employed as followings shown. First, the validation error was calculated and monitored after each epoch during training. In our settings, the learning rate would be reduced with rate 0.1 after the validation error stagnated for 6 epochs. Besides training would be terminated if validation error stagnated for 12 epochs. Second, to the ALL and HEM cells discrimination, since the false negative predictions generally cause higher risk than false positive ones in clinical situations, herein, in order to reduce the false negative rate, different class weights were applied to ALL cells and HEM cells empirically when optimizing loss function (Eq. 1) of ‘combined’ model (*λ*_All_ = 2.5 for ALL cells and *λ*_Hem_ = 1.0 for HEM cells).

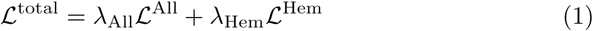

All the training and validation operations were performed on one Ubuntu 16.04 server (with one GPU 1080), employing Keras (ver. 2.2.4) with Tensorflow (ver. 1.12.0) backend.

### 2.6 Inference

The final predictions were made using the combinations of Model A, B and ‘combined’ one. Specifically, the output prediction probabilities of each model for one sample were compared together and the label of the prediction which had the maximum probability was employed as the final result.

## 3 Results

Fig. 4 shows the loss and accuracy variations on one training subset during training. Note that the training phase terminated (around 20 epoch) when the criterion described in Sec. 2.5 met for reducing the over-fitting of the model.

**Fig. 4.**
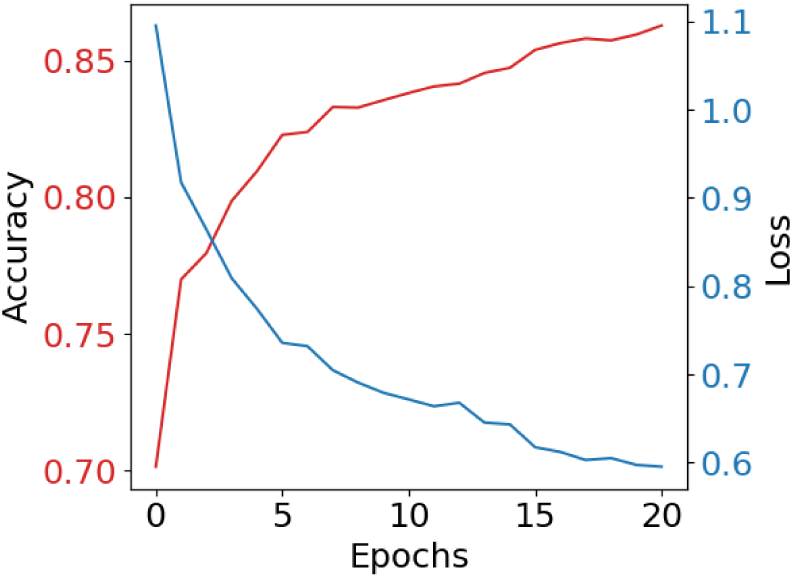
Accuracy and loss of training subset during training phase

Precision, recall and *F*_1_-score were employed as required by the contest to evaluate the performance of the model. Table 1 lists the results for the preliminary test set.

**Table 1.** Model performances of preliminary test set

| Cell type | Precision | Recall | $F_1$-score | Support |
| --- | --- | --- | --- | --- |
| Hem | 0.80 | 0.74 | 0.77 | 648 |
| All | 0.87 | 0.90 | 0.88 | 1219 |
| weighted avg. | 0.84 | 0.85 | 0.84 | 1867 |

Due to the nature of imbalance preliminary test set, weighted average (support-weighted mean per label) was also reported in Table 1. From the table, it is clear that the average prediction accuracy of ALL cells is higher than the one of HEM cells, which from the real clinical applications is more acceptable since mistakenly classifications of HEM cells to ALL cells suffer much lower risk than reversed predictions (ALL cells to HEM cells). The *F*_1_-score of final test set is 0.876.

## 4 Discussions and conclusion

In the manuscript, we proposed a two-stage image augmented enhanced bagging ensemble training scheme to solve the classification of ALL cells and HEM cells. From the experiments, the training strategy, as well as elaborately designed training subsets, are efficient for tackling the challenges (deficiency training samples and fine-grained image classification). One point needs to be noticed is that through our training parameters settings, less ALL cells prediction error is made compared with HEM cells ones. Also we would like to point out that although utilizing the model ensemble strategy to increase the overall performance, our solution is lighter compared with the one employing multiple deep models to boost the accuracy, which are quite impractical in real situations due to higher model loading burden and longer prediction latency.

In the future, more training samples are necessary for further increase the overall performance of model predictions. Besides, the blood cells detection from the scanned images should be included to make the whole procedure pragmatic. Finally, the efficiency of DL based ALL cell image analysis has to be validated with real clinical environments, not this kind of well-defined contest.

## References

1. Disease, G.., Incidence, I., Collaborators, P.: Global, regional, and national incidence, prevalence, and years lived with disability for 310 diseases and injuries, 19902015: a systematic analysis for the Global Burden of Disease Study 2015. Lancet (London, England) 388(10053), 1545–1602 (Oct 2016). https://doi.org/10.1016/S0140-6736(16)31678-6, https://www.ncbi.nlm.nih.gov/pmc/articles/PMC5055577/

2. Duggal, R., Gupta, A., Gupta, R.: Segmentation of overlapping/touching white blood cell nuclei using artificial neural networks. In: CME Series on Hemato-Oncopathology, All India Institute of Medical Sciences (AIIMS). New Delhi, India (2016)

3. Duggal, R., Gupta, A., Gupta, R., Mallick, P.: SD-Layer: Stain Deconvolutional Layer for CNNs in Medical Microscopic Imaging. In: Descoteaux, M., Maier-Hein, L., Franz, A., Jannin, P., Collins, D.L., Duchesne, S. (eds.) Medical Image Computing and Computer Assisted Intervention MICCAI 2017. pp. 435–443. Lecture Notes in Computer Science, Springer International Publishing (2017)

4. Duggal, R., Gupta, A., Gupta, R., Wadhwa, M., Ahuja, C.: Overlapping Cell Nuclei Segmentation in Microscopic Images Using Deep Belief Networks. In: Proceedings of the Tenth Indian Conference on Computer Vision, Graphics and Image Processing. pp. 82:1–82:8. ICVGIP’16, ACM, Guwahati, Assam, India (2016). https://doi.org/10.1145/3009977.3010043, http://doi.acm.org/10.1145/3009977.3010043, event-place: Guwahati, Assam, India

5. Gupta, R., Mallick, P., Duggal, R., Gupta, A., Sharma, O.: Stain Color Normalization and Segmentation of Plasma Cells in Microscopic Images as a Prelude to Development of Computer Assisted Automated Disease Diagnostic Tool in Multiple Myeloma. Clinical Lymphoma, Myeloma and Leukemia 17(1), e99 (Feb 2017). https://doi.org/10.1016/j.clml.2017.03.178, https://www.clinical-lymphoma-myeloma-leukemia.com/article/S2152-2650(17)30468-8/abstract

6. Li, C.: Classifying imbalanced data using a bagging ensemble variation (BEV). In: Proceedings of the 45th Annual Southeast Regional Conference. pp. 203–208. ACM-SE 45, ACM (2007). https://doi.org/10.1145/1233341.1233378, http://doi.acm.org/10.1145/1233341.1233378

7. Rehman, A., Abbas, N., Saba, T., Rahman, S.I.U., Mehmood, Z., Koli-vand, H.: Classification of acute lymphoblastic leukemia using deep learning. Microscopy Research and Technique 81(11), 1310–1317 (Nov 2018). https://doi.org/10.1002/jemt.23139

8. Ruder, S.: An overview of gradient descent optimization algorithms. arXiv:1609.04747 [cs] (Sep 2016), http://arxiv.org/abs/1609.04747, arXiv: 1609.04747

9. Shafique, S., Tehsin, S.: Acute Lymphoblastic Leukemia Detection and Classification of Its Subtypes Using Pretrained Deep Convolutional Neural Networks. Technology in Cancer Research & Treatment 17 (Sep 2018). https://doi.org/10.1177/1533033818802789, https://www.ncbi.nlm.nih.gov/pmc/articles/PMC6161200/

10. Szegedy, C., Ioffe, S., Vanhoucke, V., Alemi, A.: Inception-v4, inception-resnet and the impact of residual connections on learning. In: AAAI Conference on Artificial Intelligence (2017)

11. Tajbakhsh, N., Shin, J.Y., Gurudu, S.R., Hurst, R.T., Kendall, C.B., Gotway, M.B., Liang, J.: Convolutional Neural Networks for Medical Image Analysis: Full Training or Fine Tuning? IEEE Transactions on Medical Imaging 35(5), 1299–1312 (May 2016). https://doi.org/10.1109/TMI.2016.2535302

12. Vununu, C., Lee, S.H., Kwon, K.R.: A Deep Feature Extraction Method for HEp-2 Cell Image Classification. Electronics 8(1), 20 (Jan 2019). https://doi.org/10.3390/electronics8010020, https://www.mdpi.com/2079-9292/8/1/20

13. Wang, Q., Wang, J., Zhou, M., Li, Q., Wang, Y.: Spectral-spatial feature-based neural network method for acute lymphoblastic leukemia cell identification via microscopic hyperspectral imaging technology. Biomedical Optics Express 8(6), 3017–3028 (May 2017). https://doi.org/10.1364/BOE.8.003017, https://www.ncbi.nlm.nih.gov/pmc/articles/PMC5480446/

14. Yu, L., Chen, H., Dou, Q., Qin, J., Wang, P.A.: Automated melanoma recognition in dermoscopy images via very deep residual networks. IEEE Transactions on Medical Imaging 36, 994–1004 (April 2017)

15. Zhang, J., Xie, Y., Wu, Q., Xia, Y.: Skin lesion classification in dermoscopy images using synergic deep learning. In: Medical Image Computing and Computer-Assisted Intervention - MICCAI 2018. vol. 11071, pp. 12–20 (2018)

